# Craving Connections: Exploring the Relationship Between Pregnancy Cravings, Maternal Health, and Child Preferences in Indian Families

**DOI:** 10.1101/2024.10.28.620189

**Authors:** M Nithya Kruthi, Shanthi Lakshmi Duraimani, Sarah Fathima, Junaid Ahmed Khan Ghori, Katherine Saikia, Annervaz KM, Balamurali AR, Rahul Ranganathan, Sabitha Thummala

**Affiliations:** Answergenomics Pvt Ltd, Delhi, India; Accenture Labs, Bangalore, India

## Abstract

Although Pregnancy cravings are a common phenomenon, their implications on maternal health, family dietary habits, and child food preferences remain underexplored, particularly in the Indian context. This study aimed to investigate the prevalence and nature of pregnancy cravings among Indian women and their potential impact on maternal health outcomes and family dietary patterns. A cross-sectional survey was conducted with 119 women who had experienced pregnancy, collecting data on craving patterns, maternal health indicators, and perceived influences on family diet. Results revealed that most women craved sugary snacks, predominantly in the second trimester. Notably, 61.9% of participants’ cravings did not align with their spouse’s food preferences, suggesting potential shifts in family dietary habits. The study also found associations between cravings, mood swings, and morning sickness, highlighting the complex interplay of physiological and psychological factors during pregnancy. These findings underscore the need for culturally sensitive, personalised prenatal care that address both the nutritional and emotional aspects of pregnancy cravings, influencing long-term family health outcomes.

## Introduction

India, with approximately 24 million births annually, is the second most populous country and ranks low on the Mother Index in Asia (Save the Children, 2015; UNICEF, 2019). Pregnancy cravings, characterised by intense desires for specific foods, are a universal phenomenon with cultural variations. While many pregnant women experience cravings, the underlying causes and potential long-term effects remain largely unexplored (Orloff, N. C. et al., 2016). Previous research has focused on the prevalence of cravings but has overlooked the dietary habits of spouses and the food preferences of children that may emerge as a result (Hook, 1978; Wijewardene et al., 1994). This study aims to address these gaps by offering insights into pregnancy cravings and their broader health and family implications within the Indian context, considering cultural, dietary, and regional influences on maternal health.

Extensive research has highlighted various aspects of pregnancy cravings, with studies indicating that up to 90% of pregnant women experience them (Orloff et al., 2016). These cravings can be categorised into sweet, salty, spicy, and sour foods, as well as non-food items in cases of pica. Investigations into the nutritional motivations behind cravings suggest they may reflect the body’s need for missing nutrients.Additionally, hormonal changes during pregnancy can alter taste and smell, while psychological factors such as stress may also play a role (Hill et al., 2016). Understanding these cravings is crucial for addressing maternal health and dietary behaviours during pregnancy

Despite the breadth of research, significant gaps remain in our understanding of their implications. Most studies have focused on cataloguing the types of foods craved and speculating on their causes, with limited attention to how these cravings affect maternal and neonatal health outcomes (Rozin et al., 1987; Tierson et al., 1985). There is a notable lack of research examining the long-term impacts of pregnancy cravings, particularly regarding future food preferences. Additionally, the influence of cravings on spouses’ dietary habits remains largely unexplored. Questions about whether these cravings lead to meaningful dietary changes within families or correlate with partners’ food preferences have yet to be adequately addressed.

The psychological aspects of pregnancy cravings require deeper exploration. Some theories suggest that cravings may be a psychological response to the stress and emotional upheaval often associated with pregnancy; however, empirical evidence supporting this idea is limited. The influence of psychological factors—such as stress, mood changes, and emotional well-being—on cravings and food consumption patterns during pregnancy remains a significant gap in our understanding. This oversight not only limits our comprehension of the complex nature of pregnancy cravings but also hinders the development of effective support and guidance for pregnant women facing intense or distressing cravings.

### Significance of Studying Pregnancy Cravings

The study of pregnancy cravings is important for both scientific and public health reasons. Understanding these cravings can offer valuable insights into maternal health and nutrition, as a woman’s dietary needs increase during pregnancy(Somer, E et al.,2002).Proper nutrition is essential for the health of both the mother and the developing foetus (Jouanne M et al .,2021). If cravings lead to an unbalanced diet, it could negatively impact maternal and neonatal health. For instance, excessive intake of sugary or fatty foods may contribute to gestational diabetes or excessive weight gain, while cravings for non-food items, known as pica, could signal underlying nutritional deficiencies that require attention(Al Nasser Y et al., 2023). Addressing these issues is crucial for ensuring the well-being of both mothers and their children.

Beyond immediate health effects, pregnancy cravings can have lasting impacts on the dietary habits of mothers and their families. Pregnant women often adjust their eating patterns to satisfy these cravings, which can influence the entire household’s dietary choices (Forbes et al., 2018). For instance, if a pregnant woman develops a craving for a specific cuisine, her partner and family members may start consuming those foods more frequently (van Lonkhuijzen et al., 2023). This shift in dietary habits can persist even after pregnancy, potentially affecting the long-term health and nutrition of the entire family.

There is increasing interest in the link between pregnancy cravings and a child’s future food preferences. Researchers suggest that the foods a woman craves and consumes during pregnancy may shape the developing foetus’s taste preferences and eating behaviours later in life (Mennella et al., 2001). For instance, if a woman frequently craves sweet or spicy foods, her child might develop a preference for those flavors. This phenomenon, known as “prenatal flavour learning,” indicates that the prenatal environment can have lasting effects on a child’s health and behaviour, making it a crucial area of study for early childhood nutrition and development (Trout et al., 2012).

The cultural significance of pregnancy cravings also warrants attention. In many societies, food is deeply intertwined with cultural identity, and pregnancy cravings can reflect this connection (McNamara et al., 2012). By studying cravings in different cultural contexts, researchers can gain a better understanding of how cultural beliefs, traditions, and practices influence maternal and child health. For example, where traditional foods and dietary practices are highly valued, understanding pregnancy cravings could provide insights into how these cultural factors interact with the biological and psychological aspects of pregnancy (Fernández-Gómez et al., 2020).

Studying pregnancy cravings can enhance public health initiatives focused on maternal and child health.By identifying common cravings and their health implications, healthcare providers can offer targeted nutritional guidance. For example, if a woman craves nutrient-poor foods, providers can recommend healthier alternatives or supplements to ensure adequate nutrition for both mother and baby (Marshall et al., 2022). Moreover, understanding the psychological and cultural aspects of cravings allows healthcare professionals to deliver holistic support that addresses physical, emotional, and cultural needs.

### Goals and Objectives of the Research

The primary goal of this research is to investigate the relationship between pregnancy cravings, maternal health, and child preferences within Indian families. Specifically, the study aims to analyse the prevalence and types of cravings among Indian women, considering variations across regions and cultural groups.

Key objectives include investigating the nutritional motivations behind cravings, assessing their reflection of nutrient deficiencies, and understanding their impact on maternal health and dietary intake. The research will also explore how pregnancy cravings influence family dietary habits and potential links to children’s future food preferences, particularly through the lens of foetal programming.

Additionally, the study will delve into the psychological aspects of cravings, focusing on emotional and cultural influences. By providing insights into these dynamics, the research aims to enhance maternal and child health through practical recommendations for healthcare providers and families to promote healthy eating habits during pregnancy.

Another important objective of this research is to assess the broader health implications of pregnancy cravings on both short-term and long-term maternal and neonatal outcomes. This includes examining the association between cravings and pregnancy-related health conditions such as gestational diabetes, hypertension, and excessive weight gain. Identifying links between cravings and these conditions will inform healthcare practices to better monitor and manage maternal nutrition, ultimately contributing to healthier pregnancy outcomes for both mothers and children.

## Data and Methods

We developed a survey to collect data on pregnancy cravings, maternal health, and related factors. Distributed through social media channels and personal networks, the survey garnered 121 responses. After excluding incomplete submissions, a final dataset of 119 responses was analysed, ensuring a robust foundation for examining the relationship between pregnancy cravings and maternal and child health outcomes.

The questionnaire was crafted based on an extensive literature review and validated through expert feedback. Experts reviewed the initial survey items, and their suggestions were incorporated into the final version to enhance relevance and accuracy, ensuring high-quality data collection.

To obtain a representative sample, we implemented a comprehensive recruitment strategy targeting pregnant women or those who had experienced pregnancy via platforms like Facebook, Instagram, and WhatsApp. A snowball sampling method (Kirchherr et al., 2018) encouraged respondents to share the survey with acquaintances. Participation was voluntary and anonymous, with consent implied through survey completion.

Data collection spanned six months, during which we sorted and reviewed completed surveys. Incomplete responses were excluded due to the sequential nature of the questions, ensuring data integrity.

The survey included multiple sections covering participants’ demographics, experiences with morning sickness, food cravings, preferences for spicy, sweet, and sour foods, as well as trimester onset of cravings. It also explored the influence of husbands’ food preferences, aversions to specific foods, and children’s preferences for foods craved during pregnancy. Participants consented to data use for research publication.

The online survey was created using Google Forms (Google, Mountain View, California) and made accessible via a QR (quick response) code. Responses were collected anonymously through social media and personal networks.

The survey data from 119 respondents provides insights into pregnancy experiences, focusing on aspects such as the number of pregnancies, symptoms like morning sickness, food cravings, and emotional dynamics, including mood swings and family dietary influences. The findings highlight the variability in maternal health, dietary preferences, and emotional responses during pregnancy.

In terms of pregnancy distribution, 53.8% (64 individuals) reported having two pregnancies, while 37% (44 individuals) experienced one pregnancy. A smaller percentage, 7.6% (9 individuals), had three pregnancies, and only 1.7% (2 individuals) reported more than three pregnancies. This indicates that most participants have gone through one or two pregnancies, reflecting a concentrated distribution within the surveyed group.

**Figure 1.**
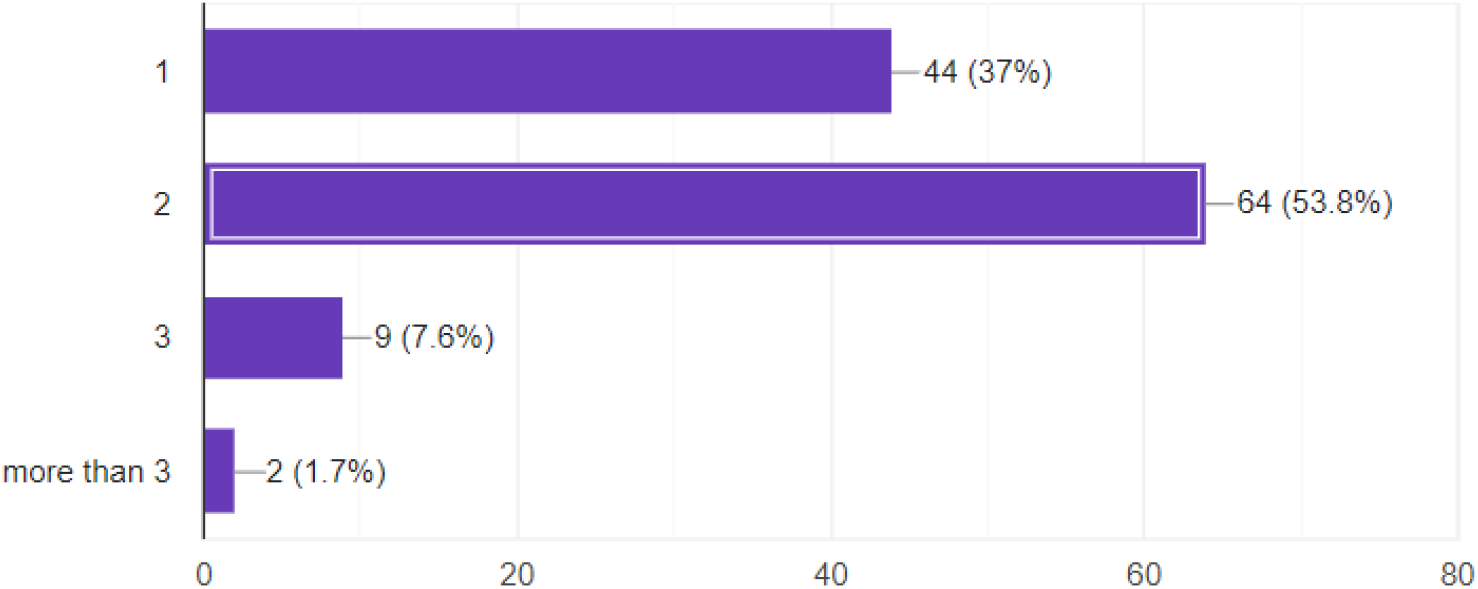
Distribution of participants based on the number of pregnancies

## Results

A majority of respondents (65.5%) experienced morning sickness during their first pregnancy, while 34.5% reported no symptoms. This indicates that morning sickness is common among pregnant women, yet the significant proportion without it highlights the diverse nature of pregnancy-related symptoms. These findings emphasise variability in maternal health experiences, suggesting that while prevalent, morning sickness is not universal. Understanding these differences can help healthcare professionals provide personalised prenatal care.

**Figure 2.**
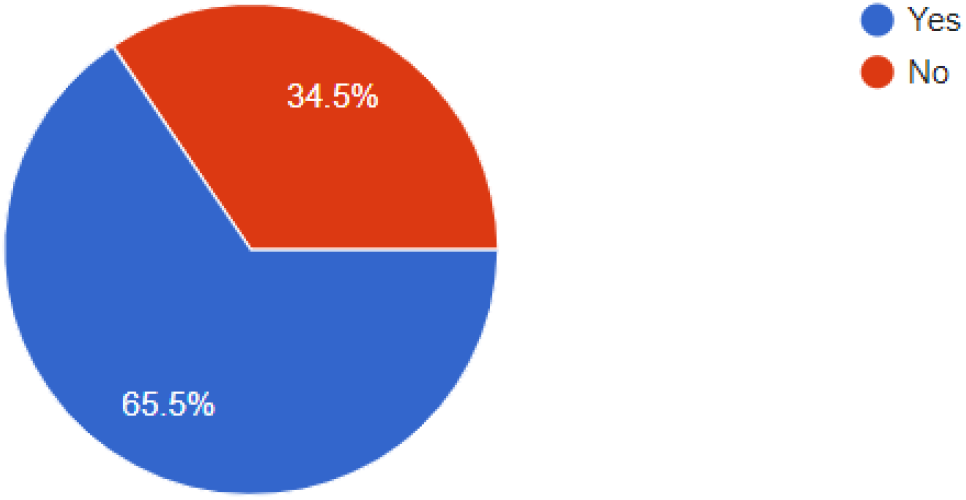
Pie chart depicting the percentage of participants who experienced morning sickness during their first pregnancy.

The onset of pregnancy cravings was most commonly reported during the second trimester, with 53% of respondents indicating this as the time cravings began. A smaller proportion, 13.3%, experienced cravings in the first trimester, while 21.7% reported cravings starting in the third trimester. Additionally, 12% experienced cravings consistently across all trimesters.

These findings suggest that cravings are more likely to emerge during the second trimester, though some women may experience them earlier or later. This variability highlights the individual nature of pregnancy experiences and underscores the importance of understanding cravings within the broader context of maternal health.

**Figure 3.**
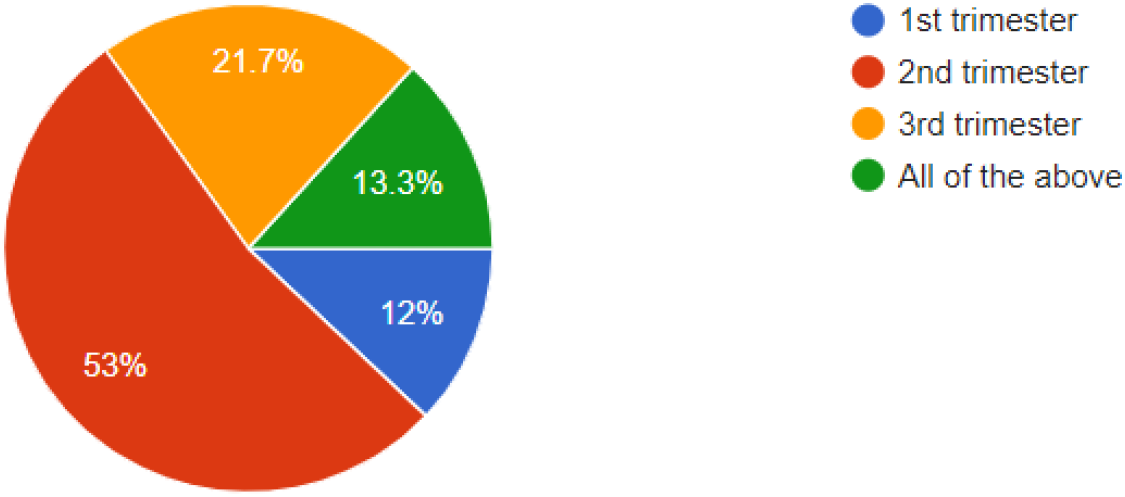
Pie chart illustrating the distribution of participants by the trimester in which they began experiencing cravings.

The survey on food cravings during pregnancy revealed diverse experiences among 83 respondents. For spicy foods, 31.3% reported cravings, 24.1% had no cravings, and 44.6% indicated that they sometimes craved spicy foods. Cravings for sweet foods were more prevalent, with 51.8% occasionally experiencing sweet cravings, 18.1% consistently having them, and 30.1% reporting no sweet cravings at all. In contrast, cravings for sour foods were less common; 21.7% of respondents reported cravings, 31.3% sometimes experienced them, and 47% had no sour food cravings. These findings highlight the varied food preferences and cravings among pregnant women, showcasing notable differences in the types of cravings experienced.

**Figure 4.**
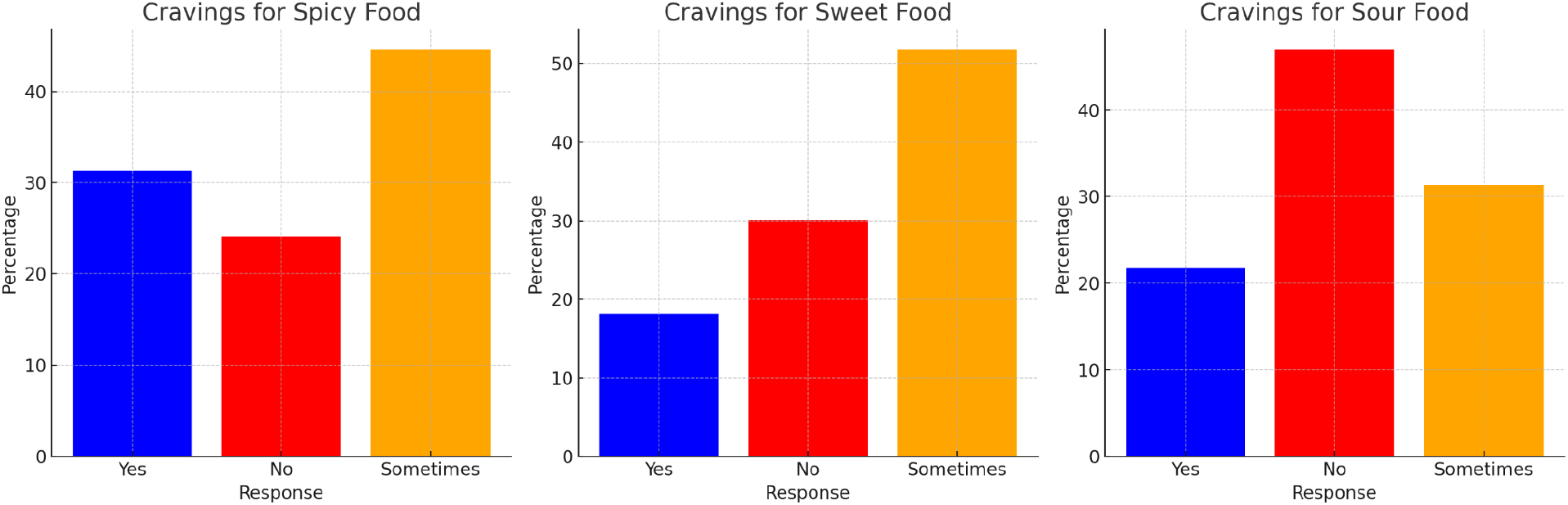
Bar graph illustrating the percentage of participants who experienced cravings for spicy, sweet, and sour foods during pregnancy.

The survey on mood swings during pregnancy found that 51.8% of respondents experienced mood swings, while 24.1% reported occasional mood swings, and another 24.1% did not experience any at all. This distribution indicates that mood swings are common for over half of the respondents, though a significant minority experienced them occasionally or not at all, reflecting variability in emotional responses during pregnancy.

**Figure 5.**
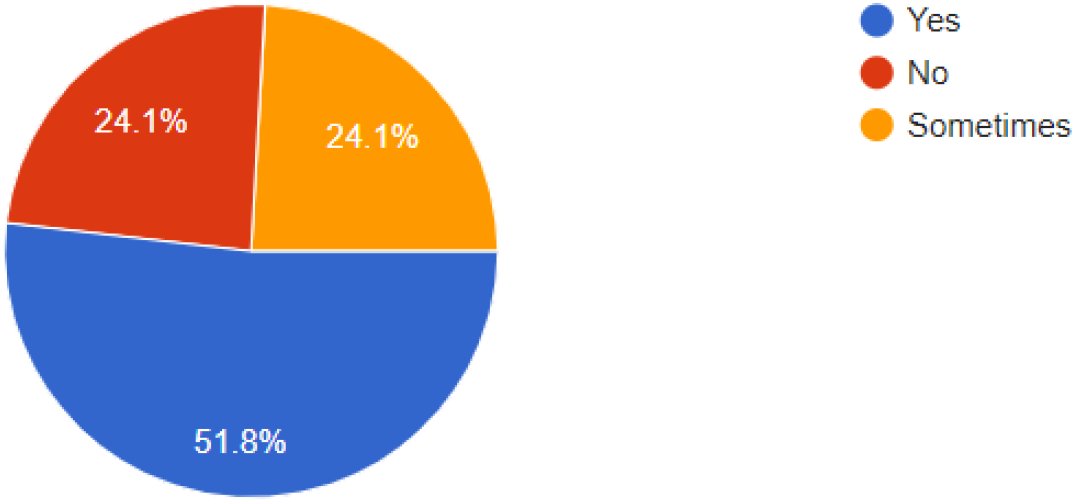
Pie chart showing the proportion of participants who reported experiencing mood swings during pregnancy.

The survey results indicate that food cravings during pregnancy often do not align with husbands’ preferences. Specifically, 61.9% of participants reported that their cravings did not coincide with their husband’s preferences. In contrast, 19% indicated that their cravings sometimes matched, and another 19% found that their cravings consistently aligned with their husband’s preferences. This suggests that while some overlap exists, most cravings appear to be independent of external influences.

**Figure 6.**
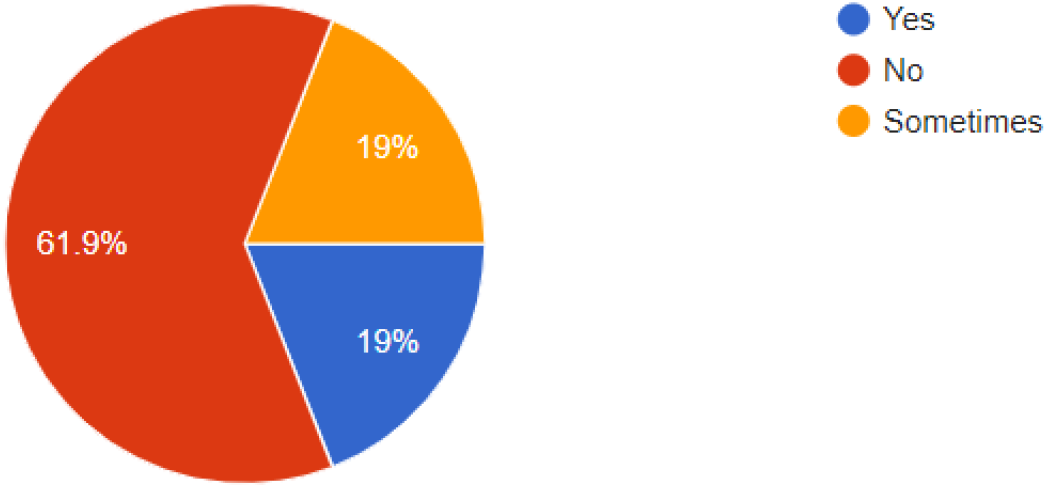
Pie chart illustrating the percentage of participants whose food cravings during pregnancy coincided with their husband’s food preferences

The results reveal that over half of respondents (52%) reported their child enjoys some of the foods they craved during pregnancy. Additionally, 27.4% indicated their child shares many food preferences, while 20.2% reported no similarities in taste. This suggests a potential connection between maternal cravings and a child’s future food preferences, although most indicated only partial overlap. These findings warrant further exploration into the long-term influence of prenatal exposure to specific foods on children’s dietary habits.

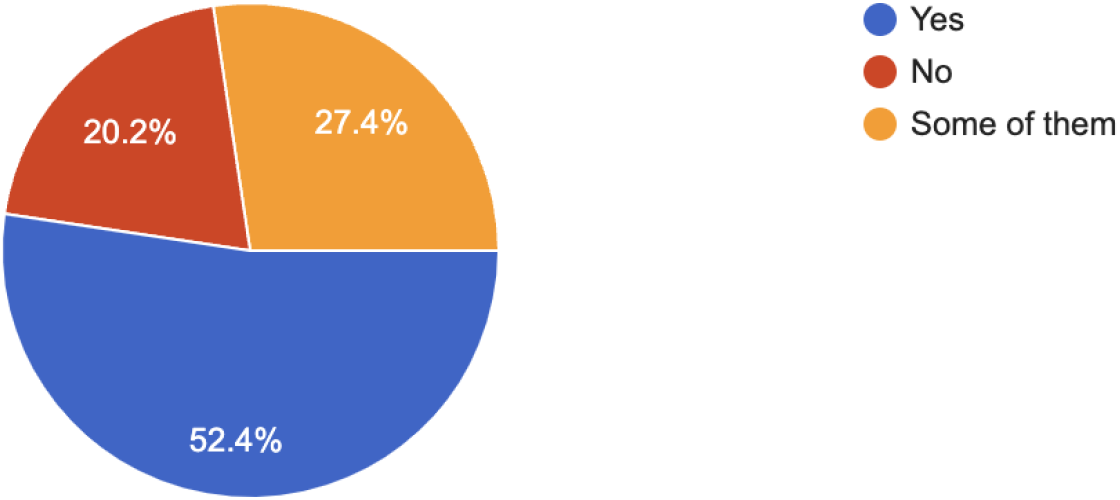

The survey results provide insights into pregnancy experiences, highlighting diversity in symptoms, cravings, and emotional responses among women. While many participants shared common experiences, such as morning sickness and cravings primarily emerging in the second trimester, the variability in these experiences underscores the individual nature of pregnancy. The findings emphasise the need for personalised care to address the unique health, dietary, and emotional needs of expectant mothers, enabling healthcare professionals to better support maternal well-being throughout pregnancy.

## Discussion

The findings from this study offer valuable insights into the variability of pregnancy cravings and their potential impact on maternal, neonatal, and family dietary habits. Consistent with previous research, most respondents reported cravings, particularly for sweet, spicy, and sour foods, which typically began in the second trimester. This variability underscores the individualised nature of pregnancy experiences and highlights the need for personalised prenatal care.

Our study confirms that pregnancy cravings are prevalent, aligning with historical research, including (Posner et al.,1957). While modern cohorts tend to favour healthier options like fruits and vegetables, the persistence of cravings emphasises their importance in maternal nutrition and cultural practices. Notably, the lower incidence of pica in the current cohort indicates a positive shift in health awareness and dietary habits over time. These findings reflect broader research showing that cravings are influenced by cultural, nutritional, and psychological factors, warranting further investigation into their long-term effects on maternal and child health.

The findings from (Hill et al.,2020) align with this research, particularly regarding the prevalence of cravings and their influence on dietary habits. Like their study, this research identifies common cravings for sweet, salty, and dairy foods among Indian women, with no significant long-term effects on maternal health outcomes. However, while Hill et al. concluded that cravings have minimal clinical impact, this study extends the investigation by exploring how cravings may influence family dietary habits and children’s future food preferences. This highlights the importance of examining both immediate and long-term effects of cravings within a familial and cultural context.

The psychological significance of cravings is supported by (Blau et al.,2020), who found that women described cravings as urgent and emotionally driven. Participants reported using strategies such as limiting food access and employing cognitive distractions to manage cravings, often feeling guilt or stress when cravings conflicted with dietary goals. This resonates with our findings, where emotional and psychological factors significantly shaped pregnancy cravings. Addressing these aspects may be crucial for promoting healthier eating habits during pregnancy and reducing the potential influence of cravings on family dietary behaviours.

Our findings reinforce the existing literature on the high prevalence of morning sickness, affecting 65.5% of respondents. This mirrors studies by (Smith et al. 2024), (Lee NM et al., 2011), and (Lacasse A et al.,2008), which indicate that nausea and vomiting during pregnancy are common experiences for many women. However, the 34.5% of participants who did not experience morning sickness highlight the variability in pregnancy-related symptoms. This variation underscores the importance of personalized prenatal care, as healthcare providers must consider the diverse experiences and symptom profiles of expectant mothers to offer tailored advice and support.

The study by (Rosen et al., 2018) highlights the multilevel determinants of maternal dietary practices, providing context for understanding the factors influencing pregnancy cravings and family dietary habits. In our research, 61.9% of women reported that their cravings did not align with their husbands’ food preferences, while 19% noted occasional or consistent alignment. This underscores the role of family dynamics, similar to the gendered food-purchasing patterns discussed by Rosen et al., in shaping maternal diets. The interaction between individual cravings and shared family meals suggests that external social influences, such as a husband’s preferences, can impact both the pregnant woman’s diet and overall household dietary patterns. This aspect of family dynamics warrants further exploration to improve maternal nutrition through both individual and family interventions.

The study by (Blau et al., 2020) offers key insights into the emotional and psychological aspects of food cravings during pregnancy, closely aligning with our findings on mood swings’ role in shaping cravings. In our research, 51.8% of respondents reported experiencing mood swings, indicating a significant emotional component to cravings. Blau et al. reinforce this connection, noting that cravings are often driven not just by nutritional needs but also as psychological responses to emotional fluctuations like stress. This supports the theory that cravings can serve as coping mechanisms, with women using food to alleviate emotional distress. Understanding these psychological drivers is crucial for developing interventions that address emotional factors, helping pregnant women manage intense cravings and reduce associated distress, thereby promoting better maternal health outcomes.

Our results reveal that 52% of participants reported their child enjoys some of the same foods they craved during pregnancy, with 27.4% indicating a strong overlap in preferences. This aligns with earlier research by (Al-Mehaisen et al., 2018), suggesting that prenatal exposure to specific foods may influence a child’s future dietary preferences. However, 20.2% of participants reported no shared preferences, indicating only a partial connection for most respondents. This suggests that while prenatal cravings may impact a child’s food choices, they are not the sole determinant. Additionally, the lack of strong association between pregnancy cravings and behavioural issues in children, as noted by Al-Mehaisen et al., implies that prenatal diet may primarily affect taste preferences rather than behaviour. These insights provide a foundation for future research on the long-term effects of prenatal food exposure, particularly regarding dietary habits and behavioural outcomes in children.

In light of the findings, it is important to explore the potential link between pregnancy cravings and underlying nutritional needs. Research suggests that certain cravings may reflect nutrient deficiencies, such as cravings for salty foods being associated with sodium needs, or sweet cravings indicating fluctuations in blood sugar levels (Hill AJ et al., 2016) (Belzer LM et al., 2010). While this study did not explicitly measure nutrient deficiencies, the prevalence of cravings for specific foods highlights the importance of understanding how these cravings might signal dietary imbalances. Addressing these potential deficiencies through targeted nutritional advice during pregnancy could improve maternal health outcomes and ensure balanced dietary intake, mitigating risks like gestational diabetes or excessive weight gain. Further research is needed to directly assess the relationship between cravings and nutrient deficiencies, as this could provide valuable insights for personalised prenatal care.

The findings of this study have important implications for maternal and child health. Understanding the diversity of cravings and their causes can help healthcare providers offer targeted nutritional advice to pregnant women. Specifically, cravings for less nutritious foods, like sweets, should be addressed with guidance on healthier alternatives to prevent complications such as gestational diabetes and excessive weight gain. Additionally, the influence of pregnancy cravings on family dietary habits suggests potential long-term effects on household nutrition.

Supporting expectant mothers in maintaining a balanced diet while accommodating cravings healthily could benefit both maternal and family health in the long term.

## Limitations of the study

One limitation of this study is its reliance on self-reported data, which may introduce recall bias regarding food cravings and dietary behaviours during pregnancy. Additionally, while the sample size is adequate for preliminary analysis, it may not fully capture the diversity of pregnancy experiences across different socio-economic and cultural backgrounds. The cross-sectional design limits conclusions about the long-term effects of pregnancy cravings on maternal and child health. Lastly, although the study addresses psychological and familial influences, more in-depth qualitative research is needed to fully understand the emotional and behavioural aspects of pregnancy cravings.

## Conclusion

This study provides insights into the prevalence and implications of pregnancy cravings among Indian women, highlighting their influence on maternal health and family dietary habits. Findings reveal diverse cravings, particularly for sweet, spicy, and sour flavours, with most occurring in the second trimester. This variability indicates that while cravings are common, they require personalised prenatal care.

The research suggests that cravings can impact not only maternal health but also family dietary patterns and children’s food preferences, aligning with the concept of foetal programming. Although further exploration is needed regarding the correlation between pregnancy cravings and children’s future preferences, the study indicates lasting effects on household nutrition.

Additionally, the emotional and psychological aspects of cravings, including mood swings, emphasise the need for holistic support systems for pregnant women that address both physical and emotional well-being. Overall, this study underscores the importance of culturally sensitive guidance and targeted nutritional advice to enhance maternal and child health, paving the way for future research into the long-term implications of pregnancy cravings and dietary habits.

## Declarations

### Ethics approval and consent to participate

The subjects have the right to agree to participate or to refuse to participate. Each subject who was willing to participate has been asked for approval (using informed consent). Every data taken was kept confidential and the subjects have the right to know the results of this study. This research follows the ethical principles of research as stated in the Declaration of Helsinki.

### Consent for publication

Not applicable

### Availability of data and materials

The datasets used and/or analysed during the current study are available from the corresponding author on reasonable request.

## Acknowledgements

Not applicable

